# Extracellular Histones Promote Calcium Phosphate-Dependent Calcification in Mouse Vascular Smooth Muscle Cells

**DOI:** 10.1101/2023.08.31.555403

**Authors:** Tomonori Hoshino, Davood Kharaghani, Shohei Kohno

**Affiliations:** Department of Calcified Tissue Biology, Hiroshima University Graduate School of Biomedical and Health Sciences, Hiroshima 734-8553, JAPAN; Neuroprotection Research Laboratories, Departments of Neurology and Radiology, Massachusetts General Hospital and Harvard Medical School, Charlestown, MA 02129, USA; Nanoscience and Advanced Materials Center, Environmental and Occupational Health Sciences Institute (EOHSI) and School of Public Health, Rutgers-New Brunswick, The State University of New Jersey, Piscataway, NJ 08854, USA; Department of Maxillofacial Anatomy and Neuroscience, Hiroshima University Graduate School of Biomedical and Health Sciences, Hiroshima, 734-8553, JAPAN

**Keywords:** chronic kidney disease (CKD), damage-associated molecular patterns (DAMPs), extracellular histones, vascular calcification, vascular smooth muscle cells (VSMCs)

## Abstract

Vascular calcification, a major risk factor for cardiovascular events, is associated with a poor prognosis in chronic kidney disease (CKD) patients. This process is often associated with the transformation of vascular smooth muscle cells (VSMCs) into cells with osteoblast-like characteristics. Damage-associated molecular patterns (DAMPs), such as extracellular histones released from damaged or dying cells, are suspected to accumulate at calcification sites. To investigate the potential involvement of DAMPs in vascular calcification, we assessed the impact of externally added histones (extracellular histones) on calcium and inorganic phosphate-induced calcification in mouse VSMCs. Our study found that extracellular histones intensified calcification. We also observed that the histones decreased the expression of VSMC marker genes, while simultaneously increasing the expression of osteoblast marker genes. Additionally, histones treated with DNase I, which degrades dsDNA, attenuated this calcification, compared with the non-treated histones, suggesting a potential involvement of dsDNA in this process. Elevated levels of dsDNA were also detected in the serum of CKD model mice, underlining its potential role in vascular calcification in CKD. Our findings suggest that extracellular histones could play a pivotal role in the vascular calcification observed in CKD.

**Graphical abstract:** 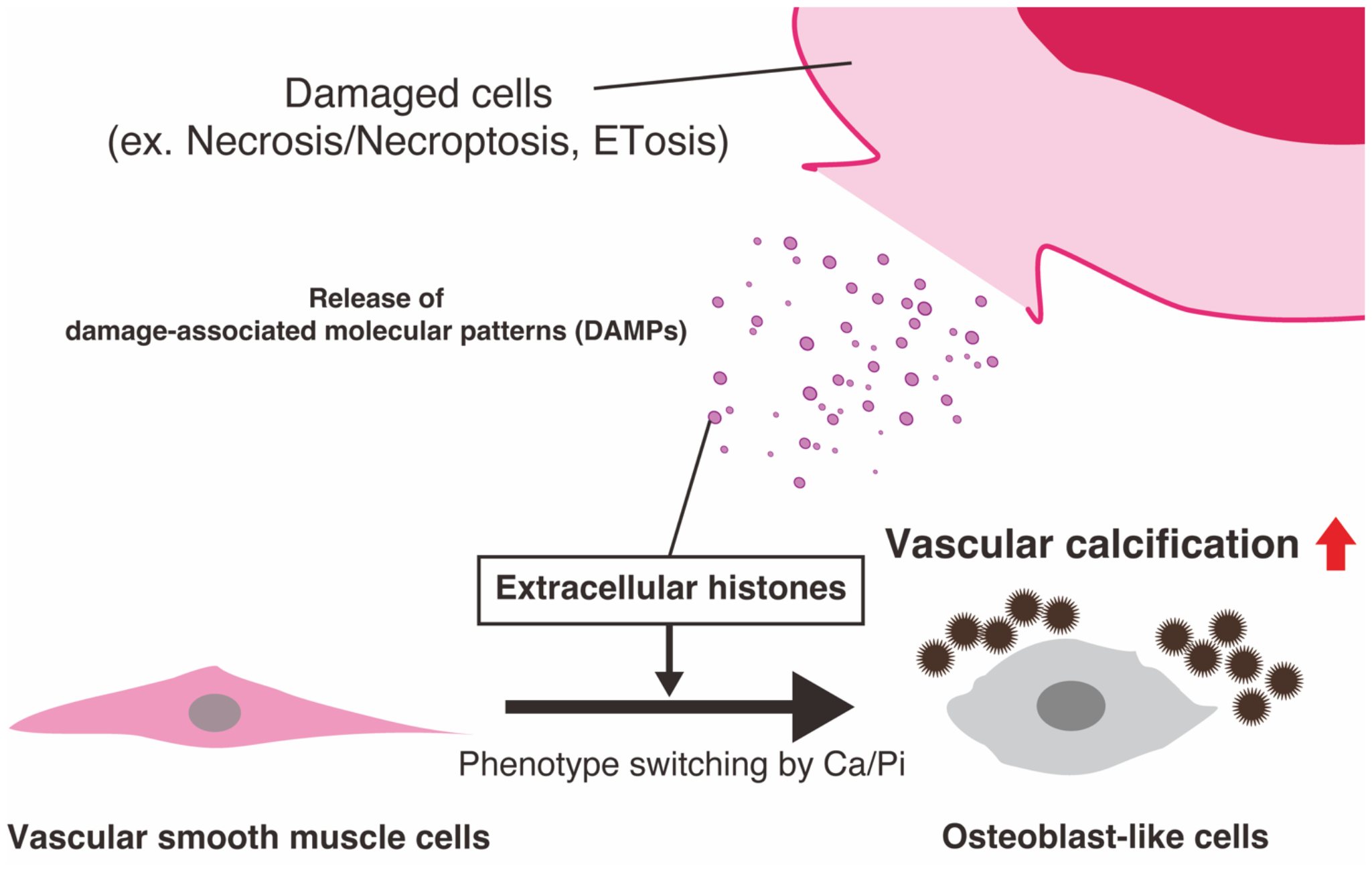

## Introduction

In patients with chronic kidney disease (CKD), vascular calcification is a significant risk factor for mortality [1]. End-stage CKD patients often have abnormal deposits of calcium in the aorta and artery (known as vascular calcification), in correlation with a phenotype switch from vascular smooth muscle cells (VMSCs) to osteogenic cells. Multiple studies have shown that the dysregulation of calcium and phosphate metabolism may be involved in the osteogenic switch of VSMCs [2], but the exact mechanism of an osteogenic switch in VSMCs is not fully understood. In addition, no treatment for vascular calcification in CKD has yet been developed.

Damage-associated molecular patterns (DAMPs), endogenous molecules released by damaged or dying cells, participate in various diseases such as cancer, neurodegenerative disease, and autoimmune disease [3]. DAMPs include a variety of intracellular components, such as histones, high mobility group box 1, heat shock proteins, and DNA (nuclear and mitochondrial) [3], and evoke sterile inflammation signaling via various receptors such as Toll-like receptors, C-type lectin receptors, NOD-like receptors, RIG-I-like receptors, and cytosolic DNA sensors [3,4].

In the aorta of CKD patients, immune cells migrate and accumulate, potentially indicating DAMPs accumulation in the lesion site [5]. Extracellular histones, as DAMPs, influence inflammation through the activation of various receptors (TLR4, TLR9, AIM2, CLEC2D) and their subsequent downstream pathways [3,4,6]. However, it remains unknown whether extracellular histones participate in vascular calcification. In this study, we evaluated the effect of extracellular histones on vascular calcification in mouse VSMCs.

## Materials and Methods

### Cell culture

Mouse VSMC cell line MOVAS (ATCC, #CRL-2797, RRID:CVCL_0F08) was cultured in high glucose Dulbecco’s Modified Eagle’s Medium (DMEM; Wako, Osaka, JPN, #043-30085) supplemented with 10% fetal bovine serum (FBS; Sigma-Aldrich, St. Louis, MO, USA, cat#173012), 100 U/mL penicillin (Sigma-Aldrich, #P3032) and 100 μg/mL streptomycin (Sigma-Aldrich, #S9137).

MOVAS cells were seeded at 20,000 cells/well in a 48-well plate (Corning, Corning, NY, USA, #353078). When they reached confluence on day 3, the cells were exposed to fresh medium supplemented with 1.8 mM CaCl_2_ (Wako, #031-00435) and 2.9 mM inorganic phosphate (Pi; NaH_2_PO_4_/Na_2_HPO_4_, pH7.4) (calcification medium) with or without histones and cultured for additional 7 days. Sterilized distilled water was used as a control. We used calf thymus-derived H2A histones (Sigma-Aldrich, #H9250; unfractionated whole histone), E. coli-derived recombinant human histone H1 (NEB, Ipswich, MA, USA #M2501), H2A (NEB, #M2502), H2B (NEB, #M2505), H3.1 (NEB, #M2503), or H4 (NEB, #M2504) (NEB’s products are proteases-free, exonucleases-free, and endonucleases-free) as extracellular histones. The cells were cultured at 37°C in a humidified atmosphere with 5% CO_2_ with medium changes every two or three days unless otherwise specified. The cells up to the seven passages were used.

### Alizarin Red S stain

After calcification induction, the cells were fixed with 4% paraformaldehyde in PBS for 20 min at room temperature, followed by staining with a 1% alizarin red S solution (200 μL/well; Sigma-Aldrich, #01-2180-2) at pH6.4, adjusted with 28% ammonia solution; Nacalai Tesque, Kyoto, JPN, #02512-95) for 30 min. The cells were washed with PBS or tap water twice at each step. Stained images were captured by BZ-X810 (Keyence, Osaka, JPN). Alizarin red S was then dissolved in 10% formic acid (250 μL/well; Wako, #066-05905), and the resultant solutions were agitated for at least 30 min, and the 100 μL of these solutions was transferred into a 96-well plate and the absorbance was measured at 450 nm using a microplate reader (Infinite M nano; TECAN, Männedorf, Switzerland). For Supplementary Fig. S1, we used the kit (PG Research, Tokyo, JPN, #ARD-SET) for Alizarin red S staining, and the dye was dissolved in 5% formic acid.

### RNA extraction and RT–qPCR analyses

Total RNA was extracted using RNAiso Plus (Takara, Shiga, JPN, #9109) or ISOGEN II (NIPPON GENE CO., Tokyo, JPN, # 311-07361) and reconstituted in RNase-free water. cDNA was synthesized from total RNA using ReverTra Ace qPCR RT Master Mix with gDNA Remover (TOYOBO, Osaka, JPN, #FSQ-301). RT-qPCR analysis was performed using the StepOne Plus Real-Time PCR system (Thermo Fisher Scientific, Waltham, MA, USA) or CFX Duet Real-Time PCR System (Bio-Rad, CA, USA) with THUNDERBIRD Next SYBR qPCR Mix (TOYOBO, #QPX-201). All processes were performed according to the manufacturer’s instructions. Expression levels for all transcripts were normalized to those of *Rpl32*. The primer sequences used for this study were as follows:

mouse *Spp1* (forward) “5′-CCATGAGATTGGCAGTGATT-3′”,

(reverse) “5′-CTCCTCTGAGCTGCCAGAAT-3′”;

mouse *Tagln* (forward) “5′-CCAGACACCGAAGCTACTCT-3′”,

(reverse) “5′-ACCCTTGTTGGCCATGTTGA-3′”;

and mouse *Rpl32* (forward) “5′-AGTTCATCAGGCACCAGTCA-3′”,

(reverse) “5′-TGTCAATGCCTCTGGGTTT-3′”,

### DNase I treatment of histones

For DNase I treatment of histones, we treated 50 μg/mL of calf thymus-derived H2A histones with 1μg/mL of DNase I (Sigma, #D4513; 2,818 U/mg) for 30 min at 37°C immediately before use. Controls were treated with the same volume of sterilized water instead of DNase I.

### CKD model mice preparation and the quantification of dsDNA

Mouse model of chronic kidney disease - mineral bone disorder (CKD-MBD) was induced by a combination of adenine diet and high phosphate [7]. Briefly, eight-to nine-week-old male C57BL/6J (CLEA Japan, Tokyo, JPN) were randomly divided into three groups (normal diet [Control], adenine diet [Adenine], Adenine diet with Phosphate diet [Adenine-Pi]). The Control group was fed the standard chow (0.8% phosphate) for 12 weeks. The adenine group and Adenine-Pi groups received 0.2% adenine for 6 weeks and then received the standard diet (0.8% phosphate) in the Adenine group or high phosphate (1.8% phosphate) in the Adenine-Pi group for 6 weeks. All studies were performed in accordance with the Guidelines for Animal Studies of Hiroshima University.

At 12 weeks after the start of the adenine diet, the mice were anesthetized with isoflurane, and the blood was collected. The blood was centrifuged at 1,500 g for 10 min at 4°C and the supernatant was collected as serum and stored at −30°C until use. The dsDNA was quantified by QuantiFluor dsDNA System (Promega, #E2671) by following the manufacturer’s instructions. Briefly, 1 μL of the serum or standard (Lamda DNA; 1ng) was mixed with 200 μL of working solution (1x QuantiFluor ON dsDNA dye diluted by TE buffer) in a 96-well plate. After 5 min of incubation, the fluorescence was measured by GloMAX (Promega) with 504 nm excitation and 531nm of emission. All samples were duplicates.

### Statistical analysis

All data are presented as mean ± standard error of the mean (SEM). Statistical differences were evaluated by either one-way analysis of variance (ANOVA) followed by Tukey’s multiple comparisons (multiple groups) or student’s t-test (two groups), using R software (Version 4.2.0). The sample size was not calculated in this study. A p-value < 0.05 was considered statistically significant.

## Results

### Extracellular histones accelerate Ca/Pi-dependent calcification in mouse vascular smooth muscle cells

To investigate the effects of extracellular histones on vascular calcification, we first evaluated whether extracellular histones (unfractionated calf thymus origin) affect calcification in MOVAS cells under high Ca/Pi conditions. As expected, extracellular histones promoted Ca/Pi-dependent calcification in a dose- and time-dependent manner (Fig. 1A and 1B; 129% and 253% increases at 10 μg/mL and 100 μg/mL, respectively, vs. Ca/Pi alone, Supplementary Fig. S1A-C) without apparent difference of morphology in the presence of extracellular histones in MOVAS cells (Supplementary Fig. S2). We also confirmed that extracellular histones enhanced Ca/Pi-dependent calcification in mouse primary VSMCs (Supplementary Fig. S3A and S3B). In addition, the elevated levels of *Spp1* (an osteoblast marker gene; Fig. 1C, 631% increase) and *Mgp* (a calcification inhibitor induced by calcification; Supplementary Fig. S4A, 123% increase) were observed, in parallel with decreased levels of VSMC marker genes such as *Tagln* (Fig. 1D, 56% decrease) and *Acta2* (Supplementary Fig. S4B, 33% decrease), when treated with extracellular histones in MOVAS cells. *Cald1*, one of the other VSMC marker, exhibited no significant difference (p = 0.1486); however, its expression level tended to decrease in response to extracellular histones (Supplementary Fig. S4C, 18% decrease). Furthermore, to investigate the impact on collagen, which plays a crucial role in calcification, we assessed the expression of genes involved in collagen production (*Col1a1* and *Col1a2*) and quantified collagen levels. However, the addition of extracellular histones showed no effect on MOVAS cells (Supplementary Fig. S5A-C).

**Fig. 1.**
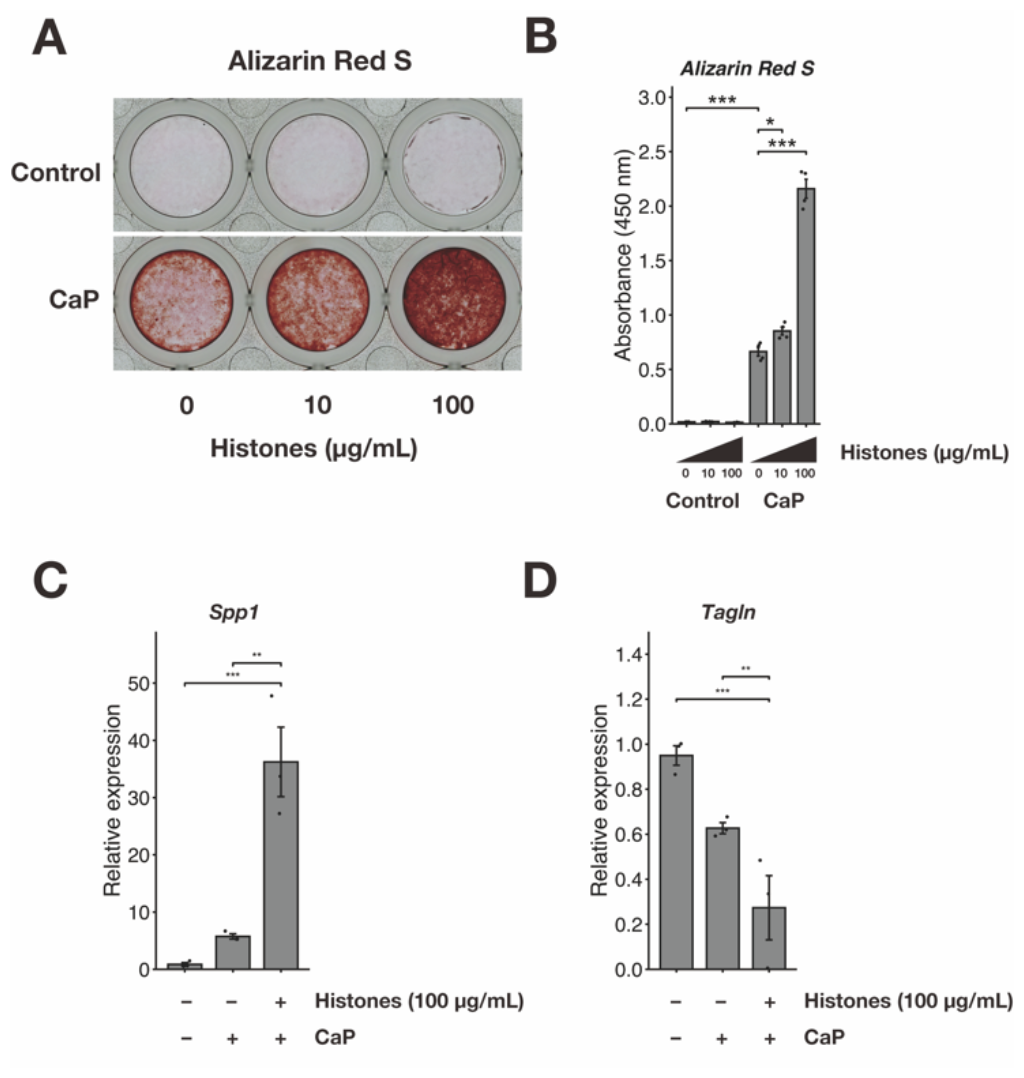
Extracellular histones accelerate Ca/Pi-dependent calcification in MOVAS cells. (**A**) The representative image of Alizarin red S stained MOVAS cells treated with calcification medium in the presence or absence of extracellular histones (0 μg/mL, 10 μg/mL, and 100 μg/mL; day 7). n = 3 per group. (**B**) Plotted to mean the dye of Alizarin red S related to (A). n = 5 per group. (**C, D**) RT–qPCR analysis of the expression of *Spp1* (**C**) and *Tagln* (**D**). The expression levels were normalized to those of *Rpl32*. n = 3 per group. All data are presented as the mean ± SEM. *: p < 0.05. **: p < 0.01. ***: p < 0.001. One-way ANOVA followed by Tukey’s post hoc test. CaP: calcium and phosphate (Ca/Pi).

Given that there are five families of histones (e.g., H1, H2A, H2B, H3.1, H4) [6], we examined the effect of human recombinant histone H1, H2A, H2B, H3.1, and H4 on Ca/Pi-dependent calcification in MOVAS cells. The results showed that different histones had different effects on Ca/Pi-dependent calcification (H1, 113%; H2A, 94%; H2B, 147%; H3.1, 466%; H4, 216% vs. Ca/Pi alone), with H3.1 being the most potent promoter of calcification in MOVAS cells under high Ca/Pi conditions (Fig. 2A and 2B). Although the molecular mechanism(s) underlying these findings are unknown, our data indicate that extracellular histones may be involved in Ca/Pi-induced calcification processes in mouse vascular smooth muscle cells.

**Fig. 2.**
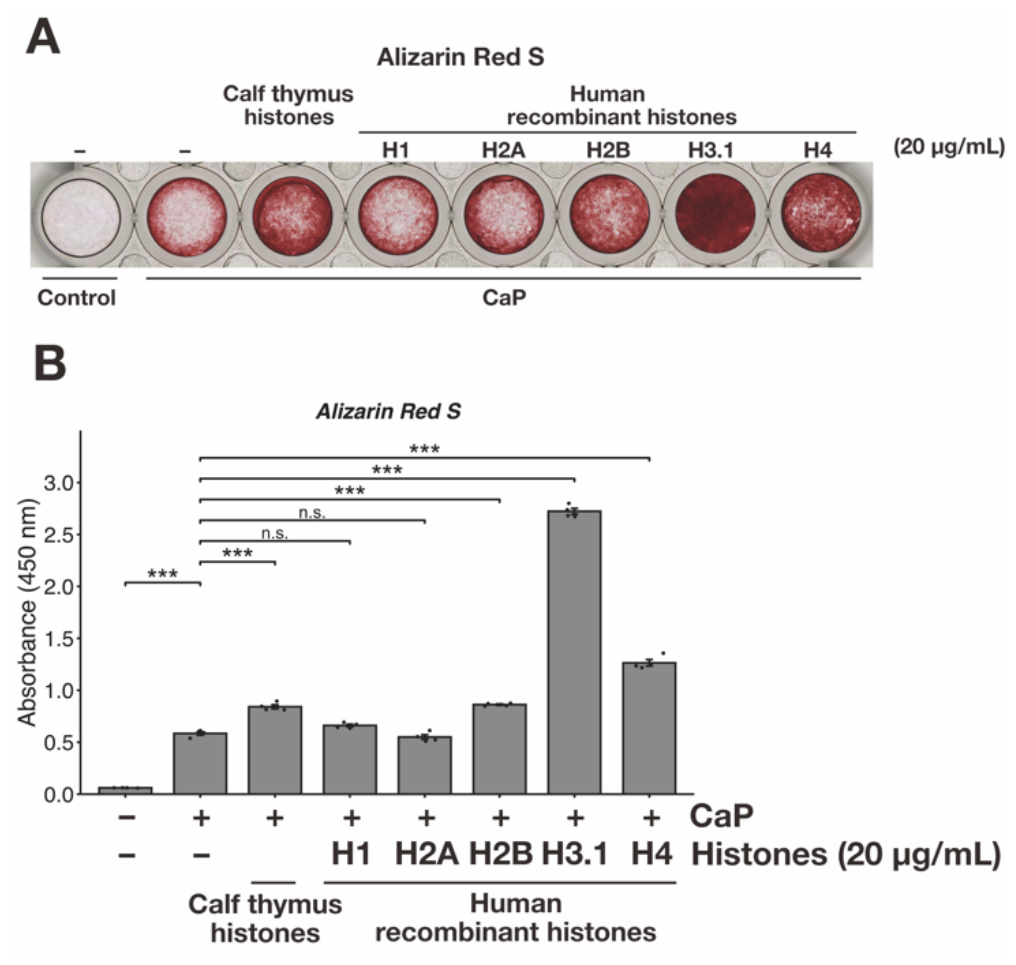
Recombinant human extracellular histones accelerate Ca/Pi-dependent calcification in MOVAS cells. (**A**) The representative image of Alizarin red S stained MOVAS cells treated with calcification medium with extracellular histones (20 μg/mL; day 7). n = 3 per group. (**B**) Plotted to mean the dye of Alizarin red S. n = 5 per group. All data are presented as the mean ± SEM. ***: p < 0.001. n.s., not significant. One-way ANOVA followed by Tukey’s post hoc test. CaP: calcium and phosphate (Ca/Pi).

Extracellular histones are known to cause cell death in certain types, such as endothelial cells, neurons, and hair follicle progenitor cells [8–11], and factors such as apoptotic bodies derived from dead cells promote vascular calcification [12]. We, therefore, performed an LDH assay (cytotoxicity assay) to assess whether extracellular histones induce cell death in MOVAS cells. We found that extracellular histones did not affect LDH leakage, i.e., cell death, in MOVAS cells, even at concentrations that promoted Ca/Pi-dependent calcification (Supplementary Fig. S6A). Because vascular calcification is known to be affected by cell density, we also examined the effect of extracellular histones on cell viability (MTT assay) in MOVAS cells. However, we could not observe any effect of extracellular histones on cell viability under normal conditions (Supplementary Fig. S6B). These results suggest that the effect of extracellular histones on Ca/Pi-induced calcification in MOVAS cells is independent of their cell death.

### DNase I-treated histones slightly attenuate Ca/Pi-dependent calcification in MOVAS cells

We further investigated how extracellular histones promote Ca/Pi-dependent calcification in MOVAS cells. Recognizing that purified histones often contain contaminating dsDNA during the purification process, we prepared DNase I-treated histones (degradation of dsDNA) and explored the potential impact of dsDNA on vascular calcification in MOVAS cells. Our findings indicate that DNase I-treated histones reduce the enhancing effect of extracellular histones on vascular calcification in MOVAS cells (Fig. 3A and 3B). In line with this observation, we detected elevated levels of dsDNA, which likely contains histones and is one of the DAMPs, in the serum of CKD model mice induced by adenine and high-phosphate diets, in which vascular calcification is highly formed (Fig. 4A and 4B), underscoring the pivotal role of dsDNA in vascular calcification.

**Fig. 3.**
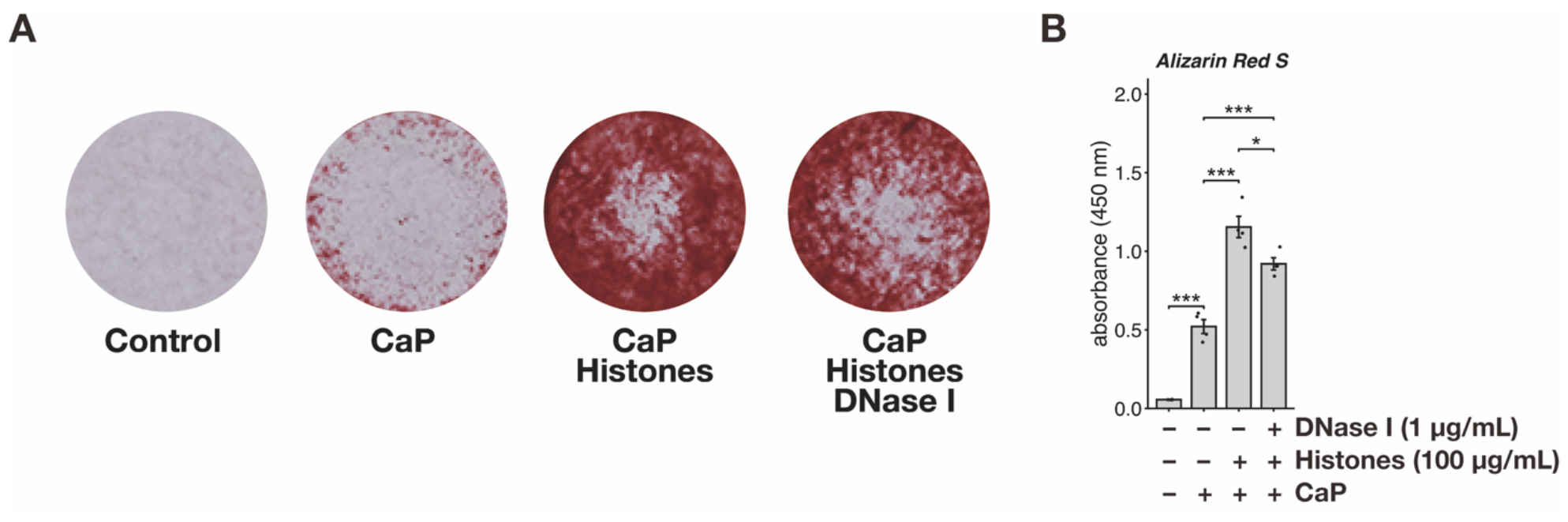
DNase I treated histones slightly attenuate Ca/Pi-dependent calcification in MOVAS cells. (**A**) The representative image of Alizarin red S stained MOVAS cells treated with calcification medium in the presence or absence of extracellular histones or DNase I treated histones (day 7). (**B**) Plotted to mean the dye of Alizarin red S. n = 3 per group. All data are presented as the mean ± SEM. *: p < 0.05. ***: p < 0.001. One-way ANOVA followed by Tukey’s post hoc test. CaP: calcium and phosphate (Ca/Pi).

**Fig. 4.**
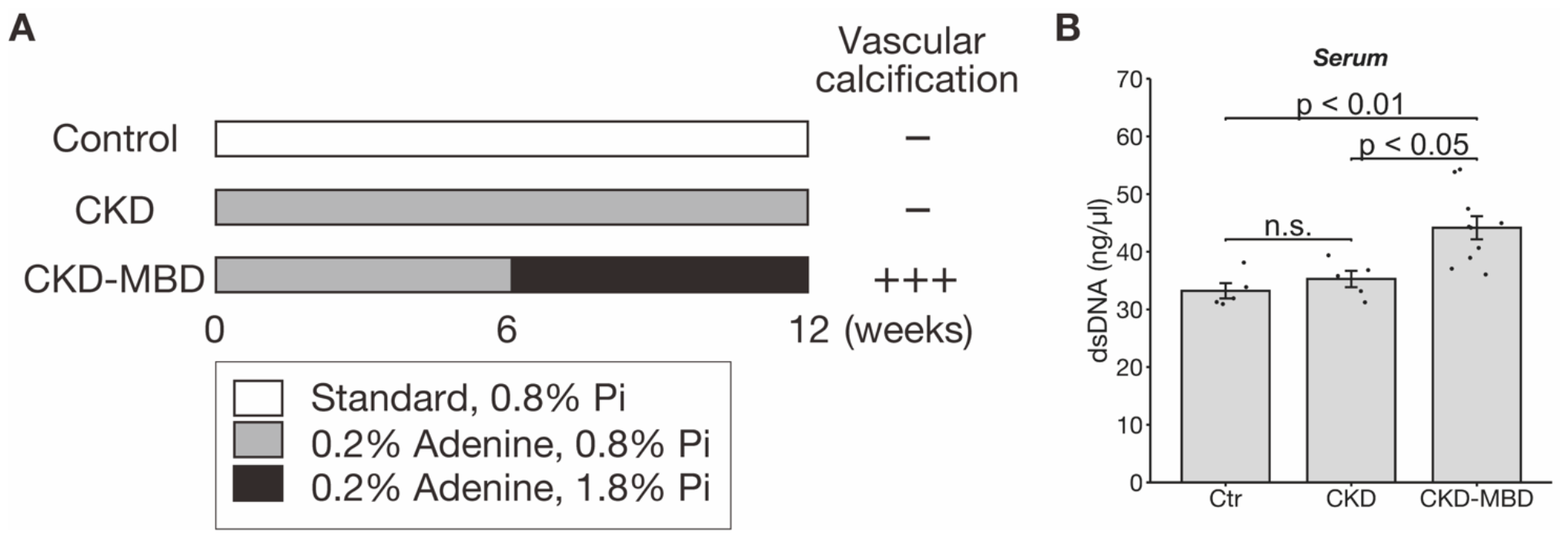
Upregulation of dsDNA levels in the serum of a mouse model of CKD-MBD. (**A**) Schematic representation of the generation of CKD and CKD-MBD mouse models. (**B**) dsDNA levels plotted to mean in the serum from control (Ctr), CKD, and CKD-MBD mice. Each group consisted of n = 5 mice for the Ctr and CKD groups and n = 10 for the CKD-MBD group. Data are presented as mean ± SEM. n.s., not significant. One-way ANOVA followed by Tukey’s post hoc test.

## Discussion

In this study, we found that extracellular histones promoted calcification in MOVAS cells (and primary mouse VSMCs) under high Ca/Pi conditions, possibly without cell death. As described, immune cells including macrophages were abundantly present in the aorta of CKD patients [5]. Given this, we reanalyzed published RNA-seq data from the aorta of an end-stage CKD mouse model (unilateral nephrectomy and high phosphorus diet model) [13]. The results showed a tendency to increase the genes of immune cell marker, such as neutrophil (*Itbg2* [CD18], *Itgam* [CD11b], *Cxcr4, Mpo*), macrophage (*Ptprc* [CD45], *Fcgr1* [CD64], *Adgre1*[F4/80], *Mertk*), eosinophil (*Ccr3, Il5ra, Siglecf*), and basophil (*Itga2b* [CD41], *Itga2* [CD49a], *Fcer1a* [FcεRI]) (Supplementary Fig. S7A-D) [13]. These results suggest that the release of DAMPs through extracellular trap cell death (ETosis) of neutrophils (NETosis), macrophages, eosinophils, and basophils and/or necrosis of the endothelial cells, VSMCs, and fibroblasts in aortic foci may be involved in vascular calcification. We also found that dsDNA associated with extracellular histones were partially involved in this calcification. Taken together with the results of the re-analysis of public RNA-seq data on the CKD mouse aorta (Supplementary Fig. S7A-D), these results suggest that the release of histones through ETosis either of neutrophils, macrophages, eosinophils, or basophils and/or necrosis either of endothelial cells, VSMCs, or fibroblasts in aortic foci may be involved in vascular calcification. Since the neutrophil/lymphocyte ratio has been reported to be increased in the blood of CKD patients [14], NETosis caused by neutrophils migrating to aortic foci may be particularly implicated in vascular calcification through the release of DAMPs. Supporting this, we have also confirmed an increase in dsDNA (probably containing histones) in the serum of CKD model mice induced by adenine and high-phosphate diets, in which highly formed vascular calcification (Fig. 4A and 4B). Thus, an increase in DAMPs, including extracellular histones and dsDNA, in aortic foci may play an important role in vascular calcification.

In this study, however, how extracellular histones promote Ca/Pi-dependent calcification in mouse VSMCs is not shown. Knockdown or knockout of the receptor of DAMPs such as TLR4, TLR9, and AIM2 suppressed vascular calcification [15–17], suggesting that the receptor of DAMPs involved in extracellular histones may be involved in the promotion of vascular calcification. In particular, TLR2, TLR4, and CLEC2D have already been reported as receptors for histones [3,4], and these receptors are involved in the induction of vascular calcification by extracellular histones. However, extracted histones often contain dsDNA. Therefore, dsDNA receptors such as AIM2 and cGAS may also be involved in their promotion of vascular calcification [3]. We have also confirmed that DNase I-treated histones ameliorate the promotion of vascular calcification by extracellular histones (Fig. 3A and 3B). However, the inhibitory effect of DNase I treatment was weak, suggesting that both histones and dsDNA may be involved in the promotion of vascular calcification. It is also possible that modifications to histones and other factors that bind to histones (proteins and RNAs such as ncRNAs) may also be involved, and these factors may interact in a complex manner. In our study, H3.1 most significantly promoted Ca/Pi-dependent calcification in MOVAS cells compared to other histone variants such as H1, H2A, H2B, and H4 (Fig. 2A and 2B). Although the detailed function is unknown, the specific structure and modifications of histones, along with the factors they interact with, could be influencing these observed effects. According to the public single-cell analysis data of adult aorta derived from mice [18], most DAMPs receptors including histones receptors (*Tlr4, Tlr7, Clec2d*) are maintained at low levels under normal conditions in VSMC cells. This suggests that extracellular histones do not directly induce transformation into osteoblasts but may enhance the process of vascular calcification. Furthermore, it has been reported that TLR4, one of the DAMPs receptors, shows increased expression upon induction of vascular calcification in cultured VSMCs [15]. This suggests that VSMCs, such as MOVAS cells, are vulnerable to DAMPs signaling, particularly when DAMPs receptor expression levels are upregulated, as seen in the induction of vascular calcification. In other words, low levels of TLR4 and other DAMPs receptors may contribute to the lack of vascular calcification in the absence of histone addition, indicating that the process of vascular calcification makes VSMCs more susceptible to DAMPs signals (Fig. 1A and 1B). Therefore, a detailed functional analysis of these receptors will further elucidate the mechanism of vascular calcification induced by DAMPs, such as extracellular histones.

Ca/Pi-dependent vascular calcification in mouse-derived VSMCs is known to be similar to that in human VSMCs (*in vitro*) and other rodent models (*ex vivo*) [19], suggesting that our findings may be linked to vascular calcification in CKD, other vascular calcification-related various diseases and aging. Thus, we expect that extracellular histones and their associated molecules may be a therapeutic target for the treatment of vascular calcification.

## Abbreviations

ANOVA: analysis of variance
CKD: chronic kidney disease
DAMPs: damage-associated molecular patterns
DMEM: Dulbecco’s Modified Eagle’s Medium
FBS: fetal bovine serum
Pi: inorganic phosphate
SEM: standard error of the mean
VSMCs: vascular smooth muscle cells

## Author Contributions

T.H. designed the experiments, performed the majority of the experiments, analyzed the data, prepared and assembled the figures, managed the project, and wrote the manuscript. S.K. assisted in performing the revised experiments. T.H. and S.K. secured funding for the research. S.K. and D.K. provided critical reading and scientific discussions. All authors reviewed and approved the final version of this manuscript.

## Acknowledgments

We thank our colleagues (S.O, R.O, T.M., and Y.Y.) at the Department of Calcified Tissue Biology, Graduate School of Biomedical and Health Sciences, Hiroshima University.

## Supplementary information

Supplementary information related to this article can be found in the online version.

