## Supplementary Information for "Extracellular Histones Promote Calcium Phosphate-Dependent Calcification in Mouse Vascular Smooth Muscle Cells"

### Supplementary Material & Methods

#### Collagen measurement

Collagen quantification was performed with the Sirius Red/Fast Green Collagen Staining Kit (Chondrex, WA, USA, #9046) according to the manufacturer's instructions. After calcification induction (day 7), the cells were washed and fixed with Kahle Fixative Solution for 10 min at room temperature. The cells were washed with PBS and stained with a Dye Solution for 30 min, followed by rinsing with distilled water repeatedly. After staining, the dye was extracted by Dye Extraction Buffer, and the eluted dye solution was transferred into a 96-well plate. The absorbance was measured at 605 nm and 504 nm using a microplate reader (Infinite M nano; TECAN, Männedorf, Switzerland).

#### Primary VSMCs culture

Primary VSMCs isolation was referred from J. of Cardiovasc. Trans. Res. (2015) with some modification [19]. Briefly, 13 weeks of age of C57BL/6J mice were euthanized by isoflurane (Pfizer, New York, NY, United States), and the aorta was carefully isolated. After perfusing the aorta with cold PBS using a tuberculin syringe (25G) to remove the blood, the aorta was placed in the DMEM with 10%FBS, 100 U/mL penicillin, 100 μg/mL streptomycin and 0.25 μg/mL fungizone (Nacalai Tesque, #02743-04). The aorta was minced until 1-2 mm piece size, and then collagenase treatment (type 2; Worthington, Columbus, OH, USA, #S8N10850) for 4-6 h. After adding the medium, the cells were centrifuged with 300g for 5 min RT, and the cells were cultured in a 24-well plate. After passaging 3 times, the cells were used for the assay.

#### Additional primer sequence

The primer sequences used for the supplementary information were as follows:

mouse *Mgp* (forward) “5′- AGGAACGCAACAAGCCTGCCTA -3′”,
(reverse) “5′- CTGCCTGAAGTAGCGGTTGTAG -3′”;
mouse *Acta2* (forward) “5′- TGCTGACAGAGGCACCACTGAA -3′”,
(reverse) “5′- CAGTTGTACGTCCAGAGGCATAG -3′”;
mouse *Cald1* (forward) “5′- GCCGTTCAAGTGCTTCACTC -3′”,
(reverse) “5′- TGTCAATCTTGGAGACTACTGCT -3′”;
mouse *Col1a1* (forward) “5′- GAGCGGAGAGTACTGGATCG -3′”,

(reverse) “5′- GTTCGGGCTGATGTACCAGT -3′”;
mouse *Col1a2* (forward) “5′- TTCTGTGGGTCCTGCTGGGAAA -3′”,
(reverse) “5′- TTGTCACCTCGGATGCCTTGAG -3′”;

**LDH assay**

The Cytotoxicity LDH Assay Kit-WST (Dojin, Kumamoto, JPN, #CK12) was used to determine cell viability, according to the manufacturer’s protocols (non-homogeneous assay). The cells were seeded at 20,000 cells/well in a 48-well plate. After 24 h incubation, the cells were then exposed to a fresh medium with or without extracellular histones for an additional 48 h. The supernatant (50 μL/well) was mixed with the same amount of the substrate solutions and incubated for 30 min at room temperature. The reaction was stopped by adding half the volume of the stop solution, and the absorbance was measured at 490 nm (Infinite M Nan, TECAN). As a positive control, MOVAS cells were treated with 20 μL/well of 10% triton-X100 (Nacalai Tesque, #12967-45) for 30 min before culture termination.

**MTT assay**

The viability (proliferation) was measured by an MTT assay. The cells were seeded at 10,000 cells/well in a 96-well plate (Corning, #353072). Next day, the cells were then exposed to a fresh medium with or without extracellular histones for an additional 48 h, followed by treatment with 5 mg/mL MTT (10 μL/well; Sigma-Aldrich, #M5655; dissolved by PBS) for 3 h before culture termination. After removing media, MTT was dissolved in 2-propanol (Wako, #166-04831) with 40 mM HCl (Wako, #080-01066), then shaken and protected from light for 10 min and absorbance was measured at 560 nm (Infinite M Nano, TECAN).

**Supplementary Figure**

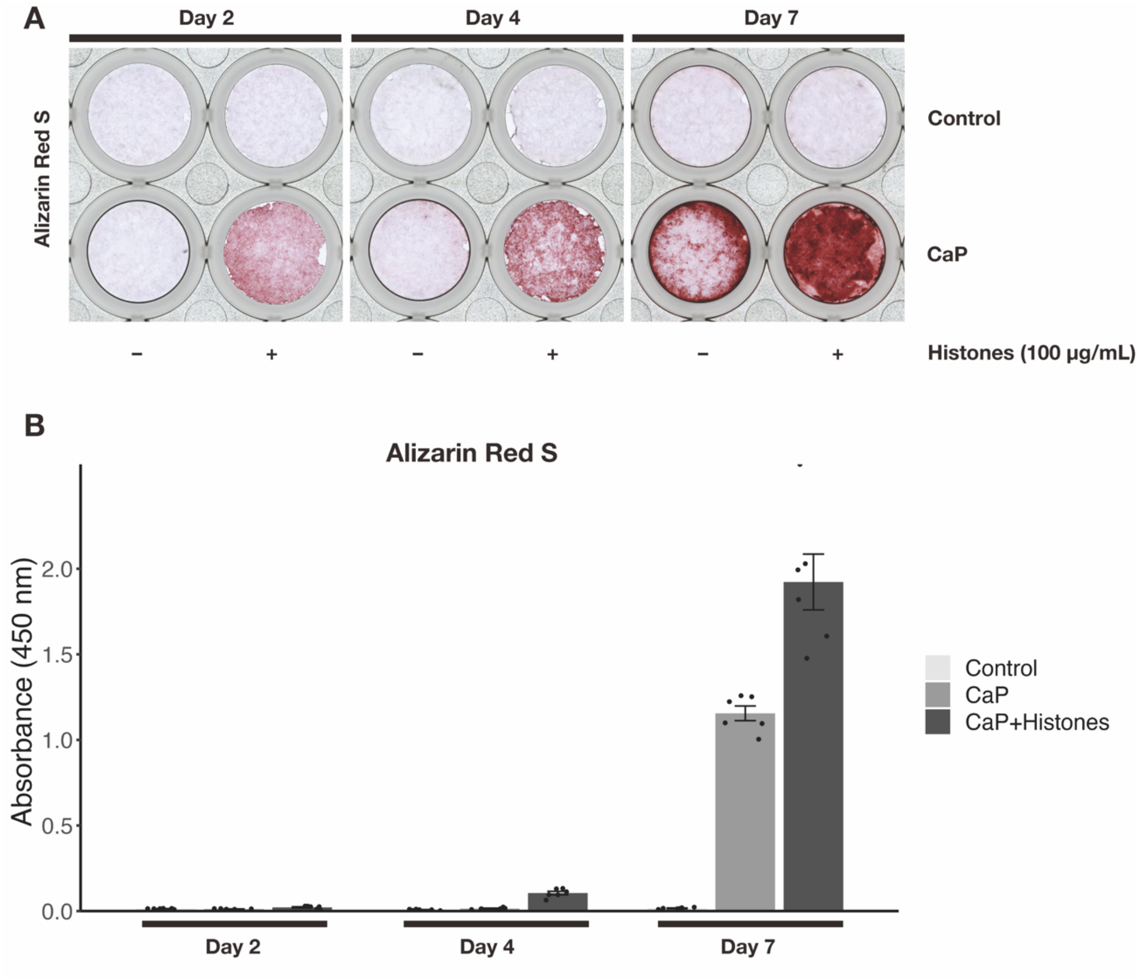


**Supplementary Fig. S1. Extracellular histones accelerate calcification formation in MOVAS cells.**

(**A**) Representative image of Alizarin red S stained MOVAS cells treated with calcification medium in the presence or absence of extracellular histones (day 2, day 4, and day 7). n = 6 per group. (**B**) The mean intensity of the dye in Alizarin red S staining related to (A). n = 6 per group. All data are presented as the mean ± SEM. CaP: calcium and phosphate (Ca/Pi).

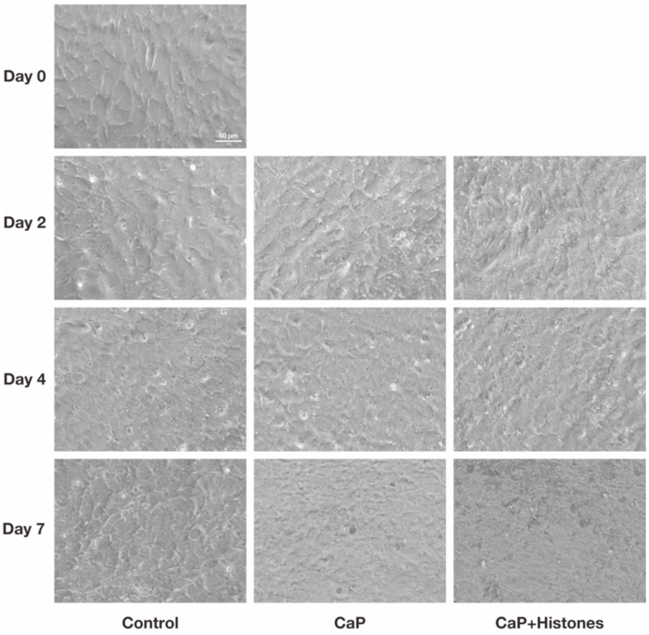


**Supplementary Fig. S2. Extracellular histones do not affect morphology in MOVAS cells.** Representative phase-contrast image of live MOVAS cells treated with calcification medium in the presence or absence of extracellular histones (day 0, day 2, day 4, and day 7). Scale bar: 50 μm.

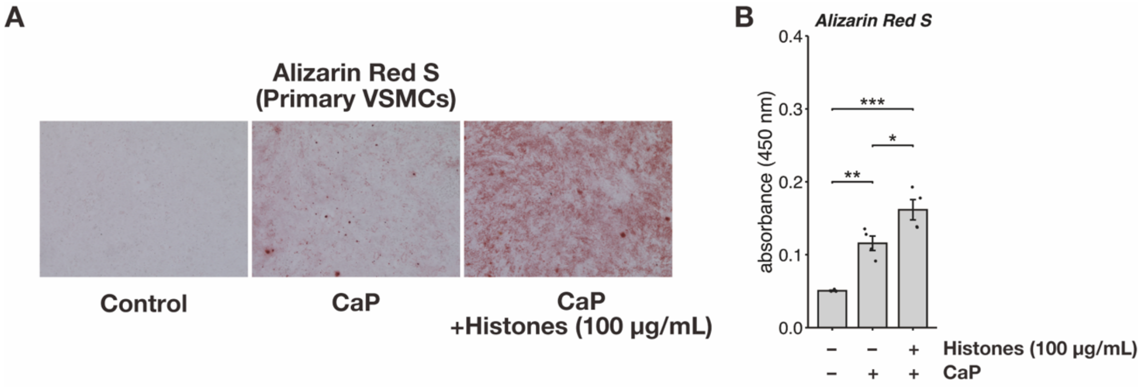


**Supplementary Fig. S3. Extracellular histones accelerate Ca/Pi-dependent calcification in primary VSMCs.**

(**A**) Representative image of Alizarin red S stained primary VSMCs treated with calcification medium in the presence or absence of extracellular histones (day 7). (**B**) The mean intensity of the dye in Alizarin red S staining related to (**A**). n = 3 per group. All data are presented as the mean ± SEM. *: $p < 0.05$. **: $p < 0.01$. ***: $p < 0.001$. One-way ANOVA followed by Tukey's post hoc test. CaP: calcium and phosphate (Ca/Pi).

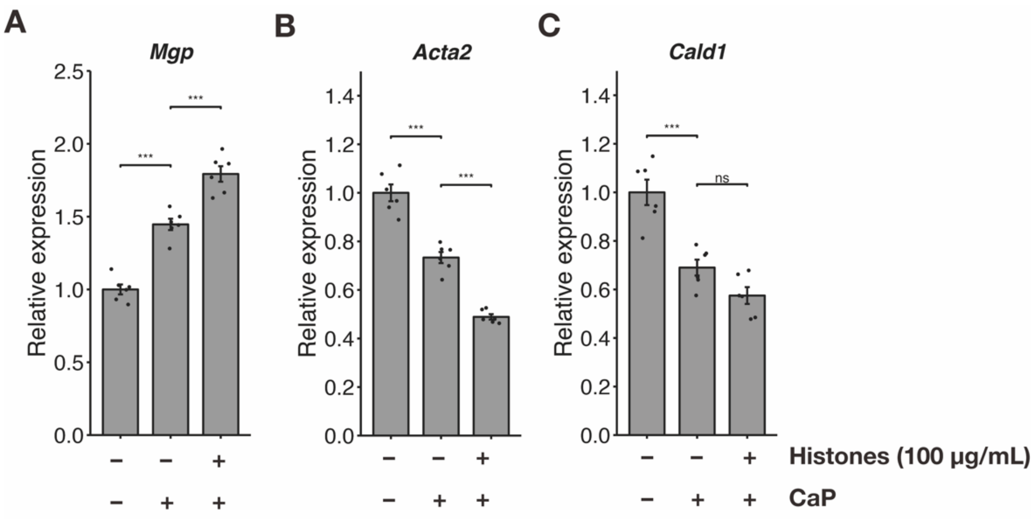


**Supplementary Fig. S4. Extracellular histones affect the expression level of *Mgp*, *Acta2*, and *Cald1* in MOVAS cells.**

(**A - C**) RT–qPCR analysis of the expression of *Mgp* (**A**), *Acta2* (**B**), and *Cald1* (**C**). The expression levels were normalized to those of *Rpl32*. n = 6 per group. All data are presented as the mean ± SEM. **: $p < 0.01$. ***: $p < 0.001$. ns: not significant. One-way ANOVA followed by Tukey's post hoc test. CaP: calcium and phosphate (Ca/Pi).

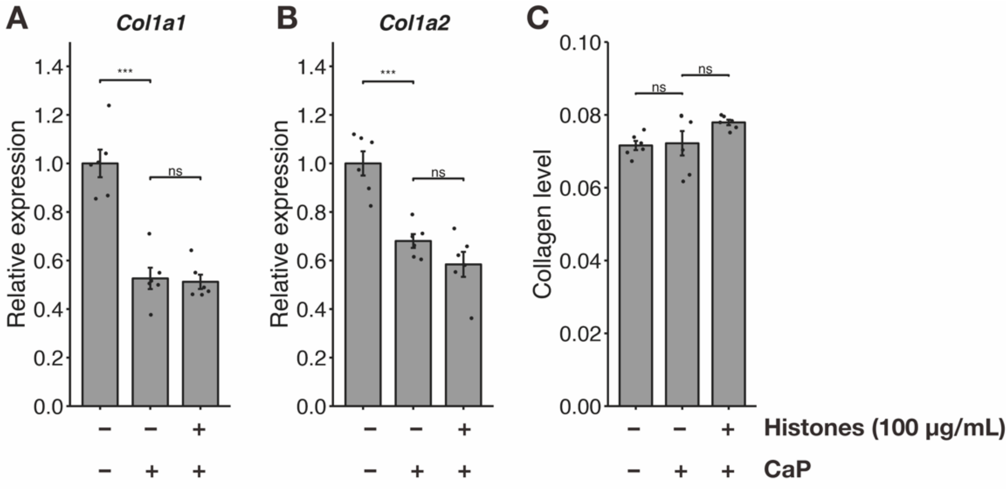


**Supplementary Fig. S5. Extracellular histones do not affect the Collagen production.**
(**A, B**) RT–qPCR analysis of the expression of *Col1a1* (**A**) and *Col1a2* (**B**). The expression levels were normalized to those of *Rpl32*. n = 6 per group. (**C**) Collagen levels in MOVAS cells treated with calcification medium in the presence or absence of extracellular histones. n = 6 per group. All data are presented as the mean ± SEM. ***: $p < 0.001$. ns: not significant. One-way ANOVA followed by Tukey's post hoc test. CaP: calcium and phosphate (Ca/Pi).

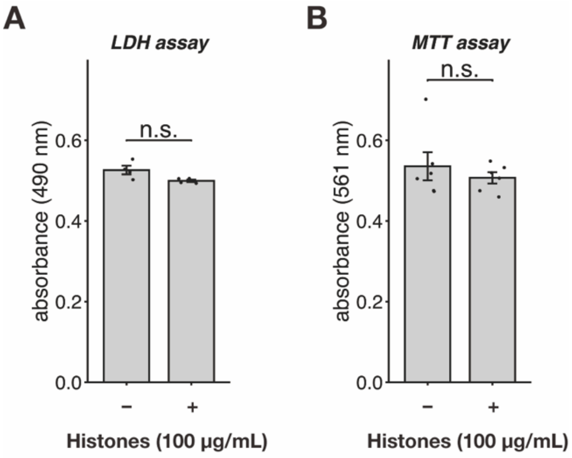


**Supplementary Fig. S6. Extracellular histones do not affect cytotoxicity and cell proliferation in MOVAS cells.**

(**A**) LDH assay (cytotoxicity) of MOVAS cells treated with extracellular histones. n = 5 per group. (**B**) MTT assay (cell proliferation) of MOVAS cells treated with extracellular histones. n = 5 per group. All data are presented as the mean ± SEM. n.s., not significant. Student's t-test.

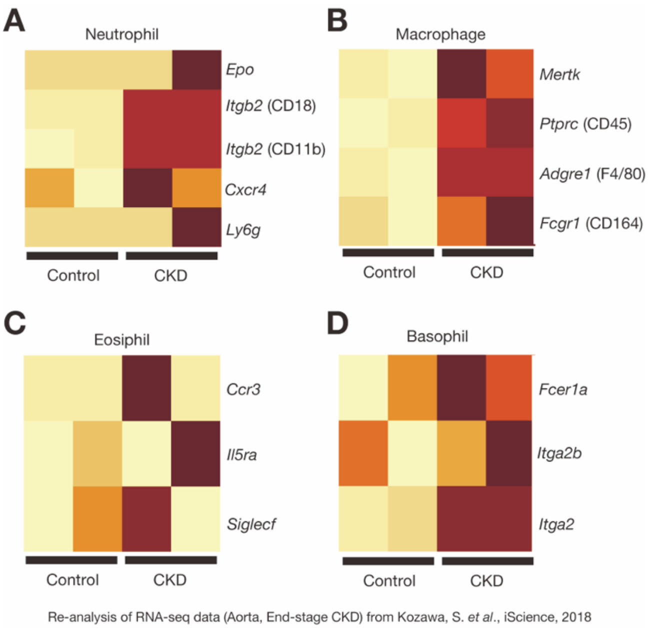


**Supplementary Fig. S7. Re-analysis of public RNA-seq data reveals the increased expression of immune cell markers in the aorta of late-stage CKD model mice.**

Re-analysis of public RNA-seq data (Kozawa S., *et al*., iScience, 2018) of the aorta of late-stage CKD model mice (Unilateral nephrectomy and high phosphorus diet model). Representative images were the heatmap that show the immune cell markers such as neutrophil (**A**), macrophage (**B**), eosinophil (**C**), and basophil (**D**). RNA-seq analyses were performed by the pipeline provided by Rhelixa, Inc. using the pipeline (FastQC v0.11.7, Trimmomatic v0.38, HISAT2 v2.1.0, Samtool v1.9, infer_experiment.py v3.0.1, featureCounts v1.6.3), and differentially expressed genes were detected by edgeR (v3.38.4) under R software (v4.2.0). We used mm10 (mouse) for the reference genome.
